# Variants in *ALDH1A2* reveal an anti-inflammatory role for retinoic acid and a new class of disease-modifying drugs in osteoarthritis

**DOI:** 10.1101/2021.09.10.457848

**Authors:** Linyi Zhu, Pragash Kamalathevan, Lada Koneva, Jadwiga Miotla Zarebska, Anastasios Chanalaris, Heba Ismail, Akira Wiberg, Michael Ng, Hayat Muhammed, Fiona E. Watt, The Oxford Hand Surgical Team, Stephen Sansom, Dominic Furniss, Matthew D. Gardiner, Tonia L. Vincent

## Abstract

Over 40% of individuals will develop osteoarthritis (OA) during their lifetime, yet there are currently no licensed disease modifying treatments for this disabling condition. Common polymorphic variants in *ALDH1A2*, that encodes the key enzyme in the synthesis of all-trans retinoic acid (atRA), have been associated with severe hand OA. In this study, we sought to elucidate the biological significance of this association. We first confirmed that *ALDH1A2* risk variants were associated with hand OA in UK Biobank. Articular cartilage was acquired from 33 consenting individuals with hand OA at the time of routine hand OA surgery. They were stratified by genotype and RNA sequencing performed. A reciprocal relationship between *ALDH1A2* mRNA and inflammatory genes was observed. Articular cartilage injury up-regulates similar inflammatory genes by a process that we have previously termed mechanoflammation, and which we believe is a primary driver of OA. Cartilage injury was also associated with a concomitant drop in atRA-dependent genes, indicative of cellular atRA levels, and both responses to injury were reversed using talarozole, a retinoic acid metabolism blocking agent (RAMBA). Suppression of mechanoflammation by talarozole was mediated by a peroxisome proliferator activated receptor (PPAR)-γ dependent mechanism. Talarozole, delivered by minipump, was able to suppress mechano-inflammatory genes in articular cartilage *in vivo* 6h after mouse knee joint destabilization, and reduced cartilage degradation and osteophyte formation after 4 weeks. These data show that boosting atRA suppresses mechanoflammation in the articular cartilage *in vitro* and *in vivo*, and identifies RAMBAs as potential disease modifying drugs in OA.

**One Sentence Summary:** Analysis of hand OA cartilage stratified by *ALDH1A2* polymorphic variants reveals a targetable, anti-inflammatory role for retinoic acid in OA.

## Introduction

Osteoarthritis (OA) is one of the leading causes of musculoskeletal morbidity, affecting all articulating joints and resulting in enormous socioeconomic burden (*1*). There are currently no disease modifying OA drugs and there is an urgent need to uncover novel treatment approaches. OA is a mechanically induced disorder whereby increased shear stress on the articulating surfaces is thought to drive protease-dependent degradation of the articular cartilage, which is associated with remodelling of the subchondral bone and synovial hypertrophy (*2, 3*). Low-grade synovitis is often observed, but anti-inflammatory strategies, such as those used in rheumatoid arthritis, have thus far failed to demonstrate clinical efficacy (*4, 5*).

We have shown that mechanical stress can activate inflammation in articular cartilage directly, through a process that we have called mechanoflammation (*6*). Indeed, injuring (cutting) the synovium or cartilage activates inflammatory signalling and induces inflammatory response genes including cyclooxygenase (*Cox2*), interleukin 6 (*Il6*), arginase-1 (*Arg1*), interleukin-1(*Il1*), C-C motif chemokine ligand 2 (*Ccl2*), and key disease related proteases such as a disintegrin and metalloproteinase with thrombospondin repeats (*Adamts4* and *Adamts5*) (*7*). The same inflammatory genes are also induced in experimental models of rodent OA in which the joint is surgically destabilized (*8–10*). Other consequences of joint stress include induction of cyclin-dependent kinase inhibitor 1A *(Cdkn1a)*, also known as p21, indicative of cell senescence, which has also been shown to contribute to OA development (*11*). The highly mechanosensitive nature of experimental OA is demonstrated by showing that after joint destabilization, inflammatory gene regulation can be abrogated by joint immobilization and this is associated with protection from developing OA (*8*).

The hands are not weight-bearing joints, but nonetheless sustain significant mechanical strains and develop OA with an overall prevalence of 27% (rising to 80% among older adults) (*12, 13*). This leads to substantial disability through pain and the loss of hand function (*14*). A genome-wide association study (GWAS) in hand OA identified the association of polymorphic variants in aldehyde dehydrogenase 1 family member A2 (*ALDH1A2*) with severe hand OA (*15*). Two of these reference single nucleotide polymorphisms (SNPs), rs3204689 and rs4238326 are common (with effect allele frequencies of 52 % and 41 % respectively) leading to overall odds ratios for severe hand OA of 1.46 and 1.44 respectively. The *ALDH1A2* gene encodes the enzyme that irreversibly catalyses the production of all-trans retinoic acid (atRA) from retinaldehyde.

atRA is an active metabolite of vitamin A, maintained at low nanomolar concentrations by a tightly orchestrated balance between its synthesis and degradation (*16*). Once delivered to the nucleus, atRA can bind to one of three retinoic acid receptors (RARα, RARβ and RARγ), or to peroxisome proliferator activated receptors (PPARβ/δ) to form a heterodimer with retinoid X receptors (RXRα, RXRβ and RXRγ). Each of these dimeric receptors binds to their respective retinoic acid response elements (RARE) or peroxisome proliferator response elements (PPRE) to activate or repress transcription of target genes (*17*). atRA isomers can also be generated and these display variable affinities at nuclear receptor binding sites (*17*). atRA metabolising enzymes, the CYP26s (CYP26A1, CYP26B1 and CYP26C1), hydrolyse atRA thus inactivating it (*18*). *RARs* and *CYP26* genes are known atRA-responsive genes and contain RAREs in their promoter regions (*19*). Blocking CYP26-mediated destruction using drugs called atRA metabolism blocking agents (RAMBAs), enhances atRA levels *in vivo* (*20–22*). Talarozole is the most specific available CYP26 inhibitor in this class (*23*).

atRA has an indispensable role in upper limb development but its role in adult cartilage is poorly understood (*19, 24–27*). Here, we interrogate the biology of atRA by performing RNA sequencing of human hand OA cartilage, stratified according to polymorphic variants in *ALDH1A2*. Using normal cartilage from mouse and pig, we reveal that atRA is a highly mechanosensitive anti-inflammatory molecule in chondrocytes, acting through PPARγ. Finally, we reveal that talarozole, by maintaining endogenous atRA levels, suppresses mechanoflammation both *in vitro* and *in vivo* and attenuates the development of post-traumatic OA. As RAMBAs have been tested clinically for other disease indications (*22, 28*), we propose that boosting cellular atRA using RAMBAs represents a novel disease-modifying strategy in OA.

## Results

### Replication of SNP associations with hand OA in UK Biobank

We first confirmed that the two biallelic genome-wide significant SNPs at the *ALDH1A2* locus (15q22), reported in the Icelandic GWAS of hand OA: rs3204689 and rs4238326 (*15*), were replicated in UK Biobank. Using primary and secondary ICD-10 diagnoses for hand OA, both SNPs were highly suggestive of association with hand OA (rs3204689 - OR 1.15 (95% CI 1.08-1.23), p = 2.5×10^−5^; rs4238326 - OR1.16 (95% CI 1.09-1.25), p = 1.1×10^−5^), being below the Bonferroni-corrected significance threshold of p<0.025 for replication of two SNPs (Table 1).

**Table 1.**
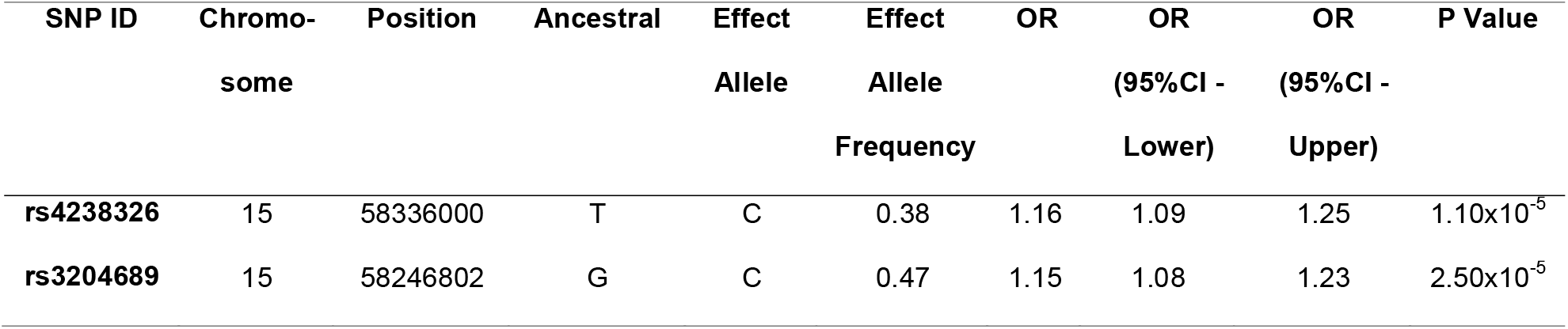
The rs4238326 and rs3204689 markers at 15q22 associated with severe hand osteoarthritis in UK Biobank cohort. Hand OA cases = 2,140, controls = 10,700 (matched on age, sex and genotyping platform).

### Polymorphic risk variants of *ALDH1A2* are associated with low levels of *ALDH1A2* in articular chondrocytes from individuals with hand OA

To explore the biological consequences of the *ALDH1A2* polymorphic variants, 33 trapezia were collected from consenting adults undergoing trapeziectomy as part of their planned usual care for symptomatic, radiographic base of thumb OA. The trapezium is the carpal bone forming the proximal part of the first carpometacarpal joint (Fig. 1A). OA at this joint exhibits classical histological features of OA (Fig. 1B). For each trapezium received, cartilage and bone were separated and snap frozen within one hour of surgery. Genomic DNA was extracted from bone for genotyping (Fig. 1C). Among the participant samples (25/33 female), we detected a high prevalence of the risk-conferring C allele in the two common SNPs in *ALDH1A2* (25/33 for rs3204689[C/G] and 22/33 for rs4238326[C/T]). The genotype and characteristics of the 33 participants are listed in supplementary Table S1. To determine the functional importance of polymorphic variants in *ALDH1A2* in hand OA, we extracted mRNA from individuals’ trapezium articular cartilage and tested the expression of *ALDH1A2* gene by real-time polymerase chain reaction (RT-PCR). *ALDH1A2* mRNA was significantly lower (HET: 0.43±0.35, HO: 0.48±0.31) in tissue from individuals who were homozygous or heterozygous for the risk variants compared with the low risk alleles (Fig. 1D). 12/13 individuals were homozygous for both alleles, consistent with their being in tight linkage disequilibrium (*15*), *ALDH1A2* levels were similar in those individuals with copies of either or both allelic risk variants.

**Fig. 1.**
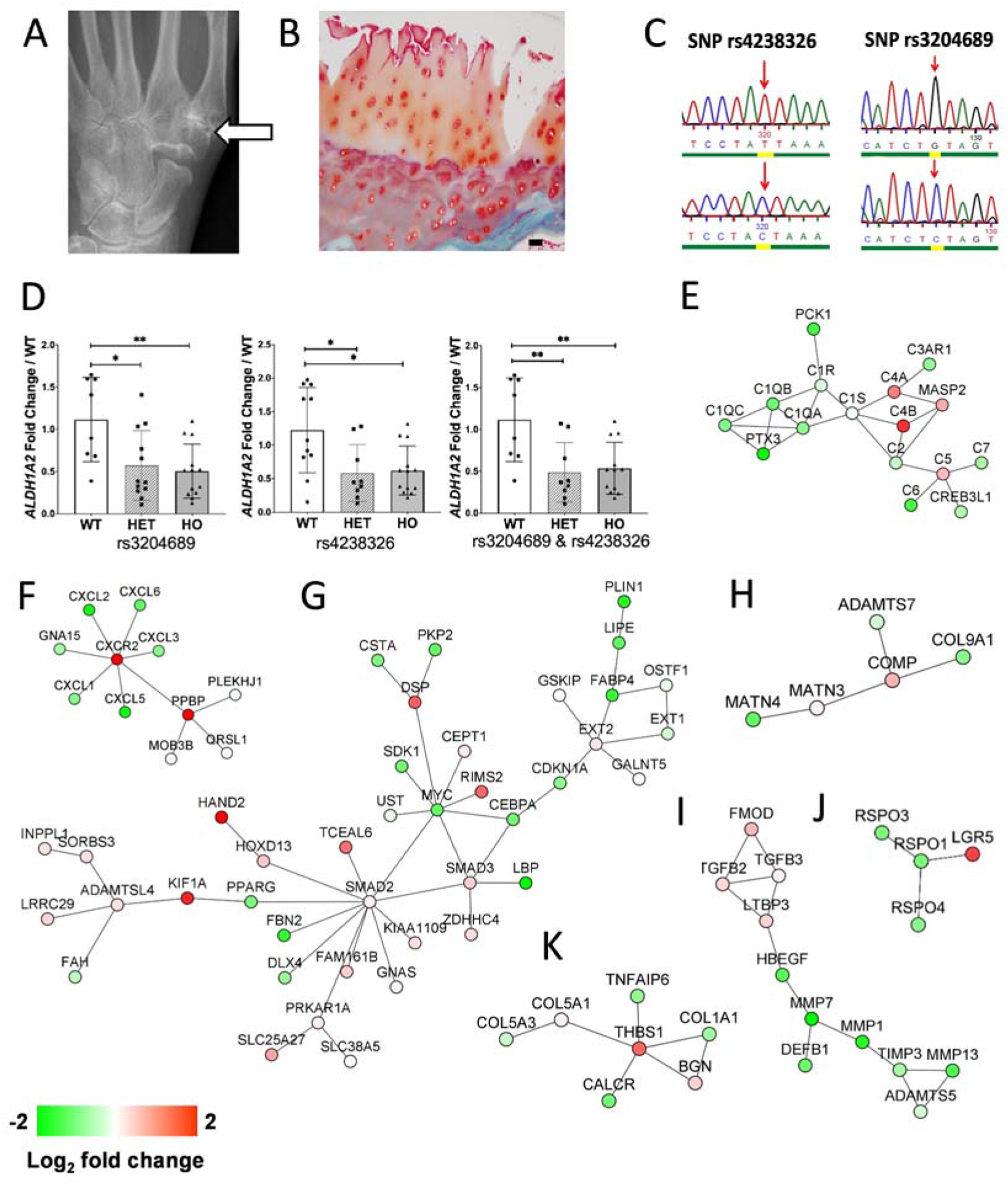
Gene regulation in cartilage from people with hand OA after stratification by polymorphic variants in *ALDH1A2*. Trapezia from 33 individuals undergoing trapeziectomy for hand OA were collected and dissected within 2h of surgery. (A) Radiographic image showing typical features of joint space narrowing, bony sclerosis and subchondral cyst formation at the first carpo-(trapezium) metacarpal joint, indicative of OA. (B) Safranin O staining of trapezial cartilage affected by OA. (C) DNA sequence analysis of two common *ALDH1A2* SNPs. (D) mRNA levels of *ALDH1A2* in wild type (WT), heterozygotes (HET) and homozygotes (HO) for rs3204689 or rs4238326 considered separately and in combination. Bars represent mean ± SEM. *p < 0.05, **p < 0.01 compared with wild type via one-way ANOVA. (E-K) Selected examples of sub-networks of osteoarthritis associated proteins that were identified by PhenomeExpress analysis as being enriched in genes that showed changes in expression level between the low and high-risk *ALDH1A2* variant patient samples: (E) complement activation, (F) chemokine-mediated signaling pathway, (G) positive regulation of osteoblast differentiation extracellular matrix organization, (H) extracellular matrix organization 1, (I) extracellular matrix organization, (J) extracellular matrix organization 2. (K) positive regulation of canonical Wnt signaling pathway chemokine-mediated signaling pathway. Nodes are colored by fold change, green: higher in HO, red: higher in WT.

### High risk *ALDH1A2* variants are associated with inflammatory gene expression pathways

To investigate the transcriptional impact of the *ALDH1A2* variants and to discover genes whose expression was associated with that of *ALDH1A2* we compared articular cartilage RNA-sequencing data from 19 patients (7 homozygous for the low-risk variants and 12 homozygous for both high-risk *ALDH1A2* variants). We first sought to identify differences in gene expression between the individuals with low and high risk *ALDH1A2* variants. Using a model that included patient gender and hand dominance as covariates, we found 15 significantly differentially expressed genes (DESeq2 analysis, BH adjusted p < 0.1) (Fig. S1 A and B, Supplementary data file S1). 10 genes with higher expression in low-risk (WT) samples included those normally expressed in erythrocytes (e.g. *HBB*, *HBA1* and *HBA2*) that showed strong expression in only a few samples in a pattern indicative of unexpected tissue heterogeneity within this group. In contrast, five genes, *CADM3*, *INO80C, RGS1, DLX4* and *FAH* showed patterns of higher expression in the samples from the patients with the high-risk *ALDH1A2* variants that were more consistent with genuine differential expression (Fig. S1B). To further explore transcriptional differences between the genotypes we adopted a pathway-level approach. Gene set enrichment analysis identified a number of Gene Ontology (GO) Biological Processes that showed higher enrichment in the high-risk (HO) samples including “response to interferon-gamma”, “regulation of IL-1 beta production” and “lymphocyte proliferation” that were suggestive of a link between the high-risk *ALDH1A2* variants and a more inflammatory phenotype (Fig. S1C and supplementary data file S2). We complemented these unbiased analyses with a PhenomeExpress analysis that was targeted to find sub-networks of proteins previously annotated to Osteoarthritis in the Human Phenotype Onotology (HP:0002758) that were enriched in genes that showed changes in expression level between the low and high-risk *ALDH1A2* variant patient samples (*29*). This analysis identified 27 sub-networks with above-background enrichments (empirical p-values < 0.05, Supplementary Table S2) including those associated with “complement activation”, “chemokine-mediated signaling pathway”, “positive regulation of osteoblast differentiation extracellular matrix organization”, “extracellular matrix organization”, “extracellular matrix organization 1”, “extracellular matrix organization 2” and “positive regulation of canonical Wnt signaling pathway chemokine-mediated signaling pathway” (Fig. 1E–K). Together, while we found relatively few genes that showed differential expression (potentially due to cohort size and heterogeneity), the pathway and network analyses provide evidence that the high-risk *ALDH1A2* variant is associated with a more inflammatory cartilage phenotype.

### *ALDH1A2* expression in hand OA cartilage is associated with reduced expression of inflammatory pathways and chemokines

We reasoned that factors other than *ALDH1A2* variant status may have influenced *ALDH1A2* expression in our patient samples. To gain insight into the biological processes associated with retinoic acid signalling in this tissue we therefore directly assessed the genes and pathways that co-varied with *ALDH1A2* expression level in osteoarthritic articular cartilage. In total, we found n=2,944 genes to be significantly positively correlated, and n=2,796 genes to be significantly negatively correlated in expression with *ALDH1A2* (Spearman correlation, BH adjusted p < 0.05, Fig. 2A and supplementary data file S3). At the pathway level, gene set enrichment analysis revealed that GO biological processes associated with higher *ALDH1A2* expression included “cilium movement” and “growth plate cartilage chondrocyte growth” while those associated with lower *ALDH1A2* expression included “regulation of cell cycle G2/M phase transition”, “osteoclast differentiation”, “Fc receptor signalling pathway”, “TNF-mediated signalling pathway” and “response to interferon-gamma” (Fig S2 and supplementary data file S4). Selected examples of significantly positively and negatively correlated genes that were also identified in the osteoarthritis-targeted PhenomeExpress sub-network analysis are shown in Fig. 2B and C. Heart and neural crest derivatives expressed 2 (*HAND2*), SMAD family member 3 (*SMAD3*), complement C4A (*C4A*), aggrecan (*ACAN*), regulating synaptic membrane exocytosis 2 (*RIMS2*), complement C4B (*C4B*), cartilage oligomeric matrix protein (*COMP*) and desmoplakin (*DSP*) positively correlated with the expression of *ALDH1A2*, while cyclin dependent kinase inhibitor 1A (*CDKN1A*), ADAM metallopeptidase with thrombospondin type 1 motif 4 (*ADAMTS4*), exostosin glycosyltransferase 1 (*EXT1*), SMAD family member 2 (*SMAD2*), matrix metallopeptidase 1 (*MMP1*), matrix metallopeptidase 13 (*MMP13*), C-X-C motif chemokine ligand 2 (*CXCL2*) and C-X-C motif chemokine ligand 5 (*CXCL5*) were inversely correlated. In keeping with the differential expression analysis results, these data suggested that lower levels of retinoic acid are associated with a more inflammatory phenotype.

**Fig. 2.**
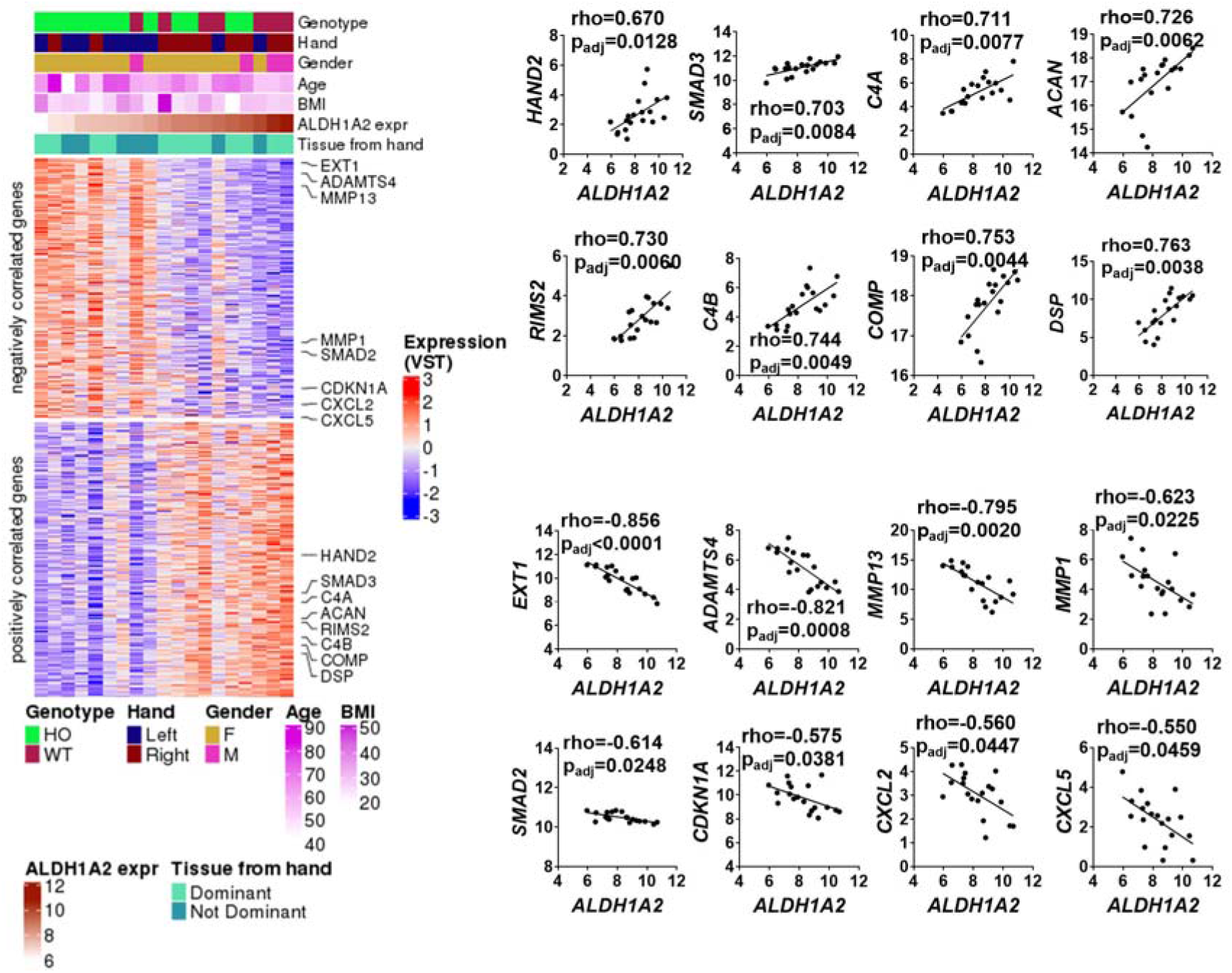
Correlation of genes with *ALDH1A2* level in hand OA cartilage. (A) Heatmap of the n=5740 genes significantly correlated with *ALDH1A2* expression (VST transformed values, BH adjusted p < 0.05). Samples (columns) are ordered according to expression of *ALDH1A2* gene in corresponding sample and genes (rows) sorted by Sperman’s coefficient correlation rho according to expression of *ALDH1A2* in data set. Participant characteristics including handedness (hand), gender, genotype and side of disease (Tissue from hand) are shown. Linear regression was performed on the central nodes identified from Phenome Express. (B) Genes that positively correlated with *ALDH1A2*. (C) Genes that negatively correlated with *ALDH1A2*.

### Cartilage injury strongly down-regulates atRA-responsive genes whilst increasing inflammatory genes

Mechanical injury is an important etiological factor in OA by triggering mechanoflammation (*9, 30*). As several genes that correlated with *ALDH1A2* were also known to be induced by cartilage mechanical injury, we investigated the effect of injury on regulation of atRA-responsive genes. atRA-responsive genes in articular chondrocytes include several genes that are part of the canonical synthetic pathway of atRA (Fig. S3). Porcine cartilage injury strongly down-regulated atRA-responsive genes *CYP26A1, CYP26B1*, and retinoic acid receptors (*RARA, RARB* and *G*) (Fig. 3A), while inflammatory genes including *CDKN1A, IL6, ADAMTS4, MMP3, TIMP1, COX2,* and *CCL2* were up-regulated as seen previously (Fig. 3B) (*31*). Similar results were seen after murine cartilage injury (Supplementary Fig. S4). Porcine cartilage injury was associated with a reduction in ALDH1A2 protein from 4h after injury (Fig. 3C, D).

**Fig. 3.**
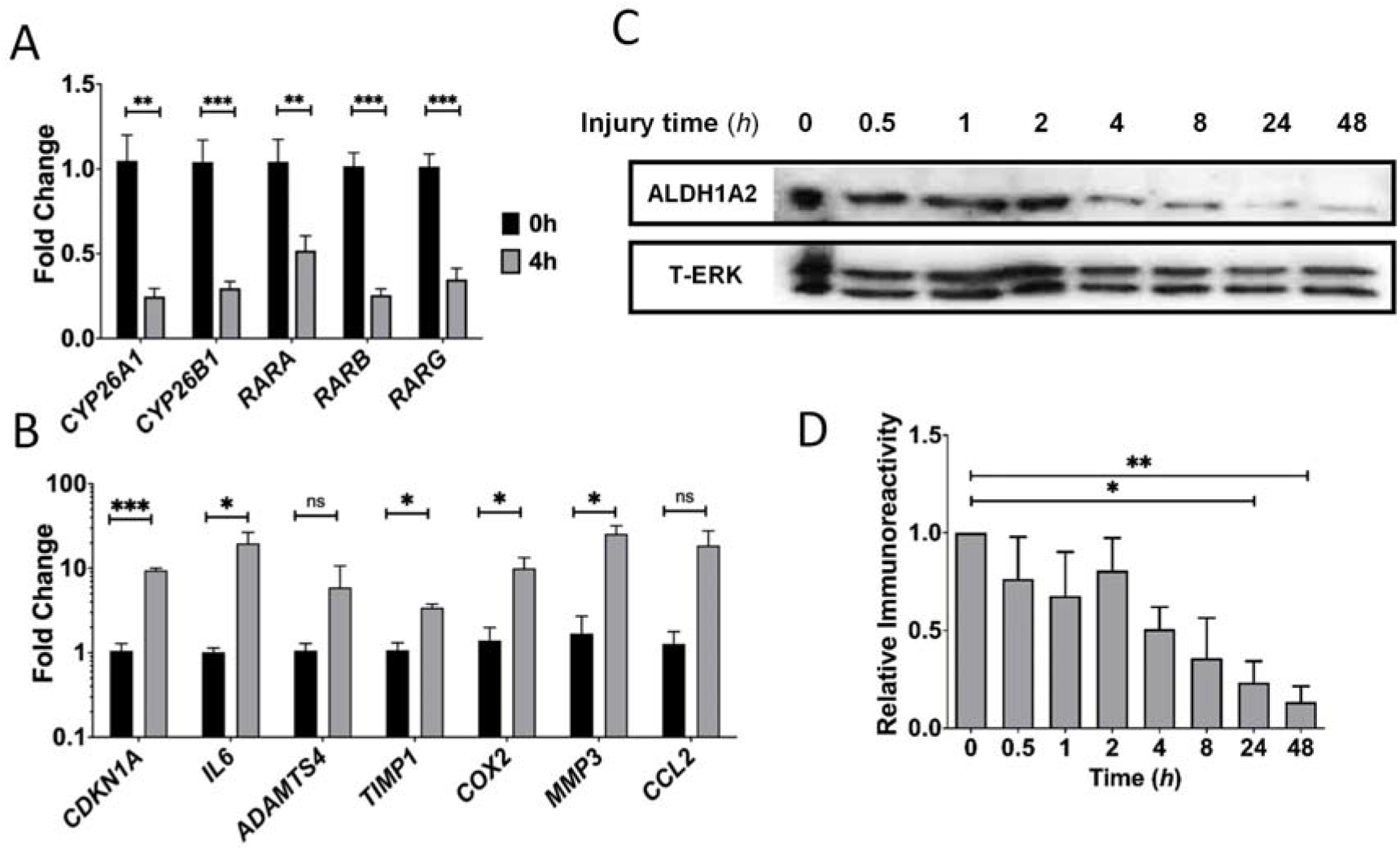
Porcine cartilage injury strongly supresses atRA-responsive genes and enhances inflammatory response genes. (A) Articular cartilage was dissected from the porcine metacarpophalangeal joint either onto dry ice (0h) or into serum free medium for 4h before extracting RNA. atRA-responsive genes (A), or inflammatory response genes (B), were measured by quantitative PCR, normalised to *18s* and expressed relative to 0h (mean ± SEM, n =3) (C) Representative ALDH1A2 protein by immunoblot from cartilage lysates post injury. (D) Quantitative analysis of ALDH1A2 protein (ALDH1A2/ERK) from three independent experiments. Statistical significance was determined using unpaired multiple t-tests. Data are mean ± SEM, n = 3 independent experiments.

### Maintaining cellular atRA levels with talarozole suppresses mechanoflammation following cartilage injury through a mechanism involving PPARγ

We next explored whether maintaining levels of atRA at time of injury could prevent mechanoflammation. Porcine metacarpophalangeal joints were pre-injected with talarozole (TLZ) or vehicle prior to cartilage injury (Fig. 4A). TLZ prevented the drop in atRA-dependent genes, *CYP26A1* and *CYP26B1*, after injury (Fig. 4B), and suppressed the regulation of several inflammatory genes following injury including *CDKN1A*, *IL6*, *ADAMTS4*, *ADAMTS5, CCL2, TIMP1*, and *COX2* (Fig. 4C). *MMP3* was the only gene significantly increased by TLZ after injury. The anti-inflammatory effect of TLZ was not specific for mechanoflammation as TLZ or atRA was able to suppress IL1-induced inflammatory gene regulation in isolated porcine chondrocytes (Fig. S5). To investigate the mechanism by which atRA, maintained by TLZ, suppressed cartilage mechanoflammation, we first examined whether TLZ was able to suppress TAK1-dependent inflammatory signalling, a known upstream mediator of mechanoflammation (*9*). Using the same porcine joint injury model (Fig. 4A), cartilage injury activated the TAK1-dependent, mitogen activated protein kinases (MAPKs), c-Jun N-terminal kinase (JNK) and extracellularly regulated kinase (ERK) within 5 minutes of cutting injury. Pre-incubation with a TAK1 inhibitor suppressed MAPK phosphorylation (Fig. 5A and B), but TLZ did not alter the pattern of MAPK activation (Fig. 5C and D), indicating that the anti-inflammatory effects of talarozole were downstream of MAPK activation. However, TAK1 inhibition did prevent the drop of atRA responsive genes indicating that this pathway is important for controlling atRA levels after injury (Fig. 5E). As PPARγ has been associated with anti-inflammatory actions (*32, 33*), we speculated that the anti-inflammatory effects of TLZ was mediated through PPREs. Injection of joints with TLZ alone or in combination with a PPARγ inhibitor (GW9662) reversed the anti-inflammatory effects of TLZ for *IL6, COX2, ADAMTS4*, and *TIMP1* indicating that PPARγ mediates the anti-inflammatory effects of TLZ (Fig. 5F). PPARγ had no effect on classical RARE-dependent genes in the presence of talarozole (*CYP26B1, RARA, RARB, RARG*) (Fig. 5G).

**Fig. 4.**
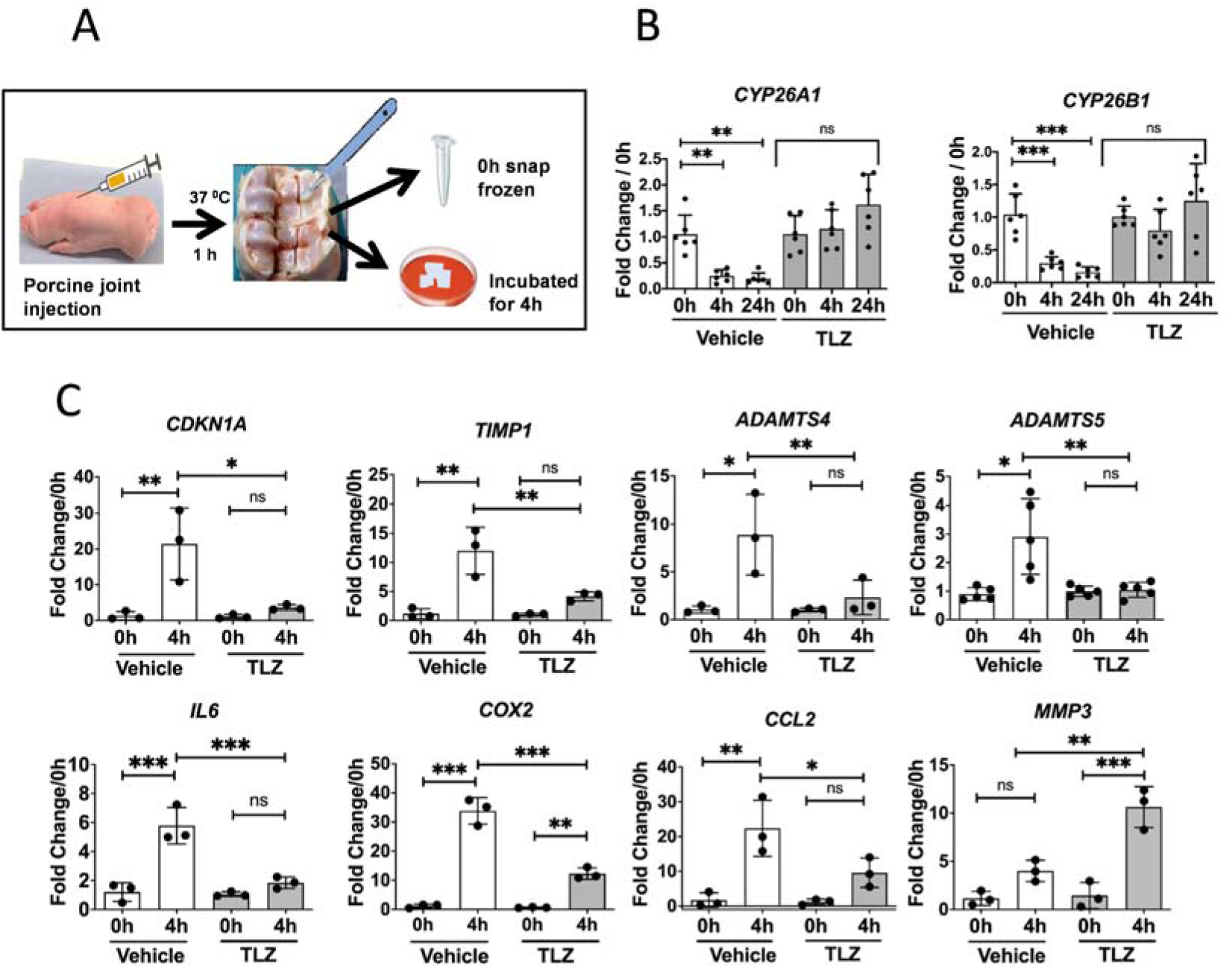
Talarozole prevents the drop of atRA-responsive genes after injury and suppresses injury-induced inflammatory genes. (A) Schematic showing the protocol for porcine MCP joint inhibitor injection. Porcine MCP joints were injected with 5 μM talarozole (TLZ) or vehicle. After 1h at 37 °C, cartilage was explanted and either snap frozen (0h) or cultured in the presence of inhibitor or vehicle for 4h. Total RNA was extracted and gene expression levels for atRA-responsive genes (B) and inflammatory genes (C), measured by quantitative PCR. n = 3-6 (as shown) for each group. Statistical analysis was performed using one-way ANOVA with Tukey’s post hoc analysis. Bars represent the mean ± SEM, *p < 0.05, **p < 0.01, and ***p < 0.001.

**Fig 5.**
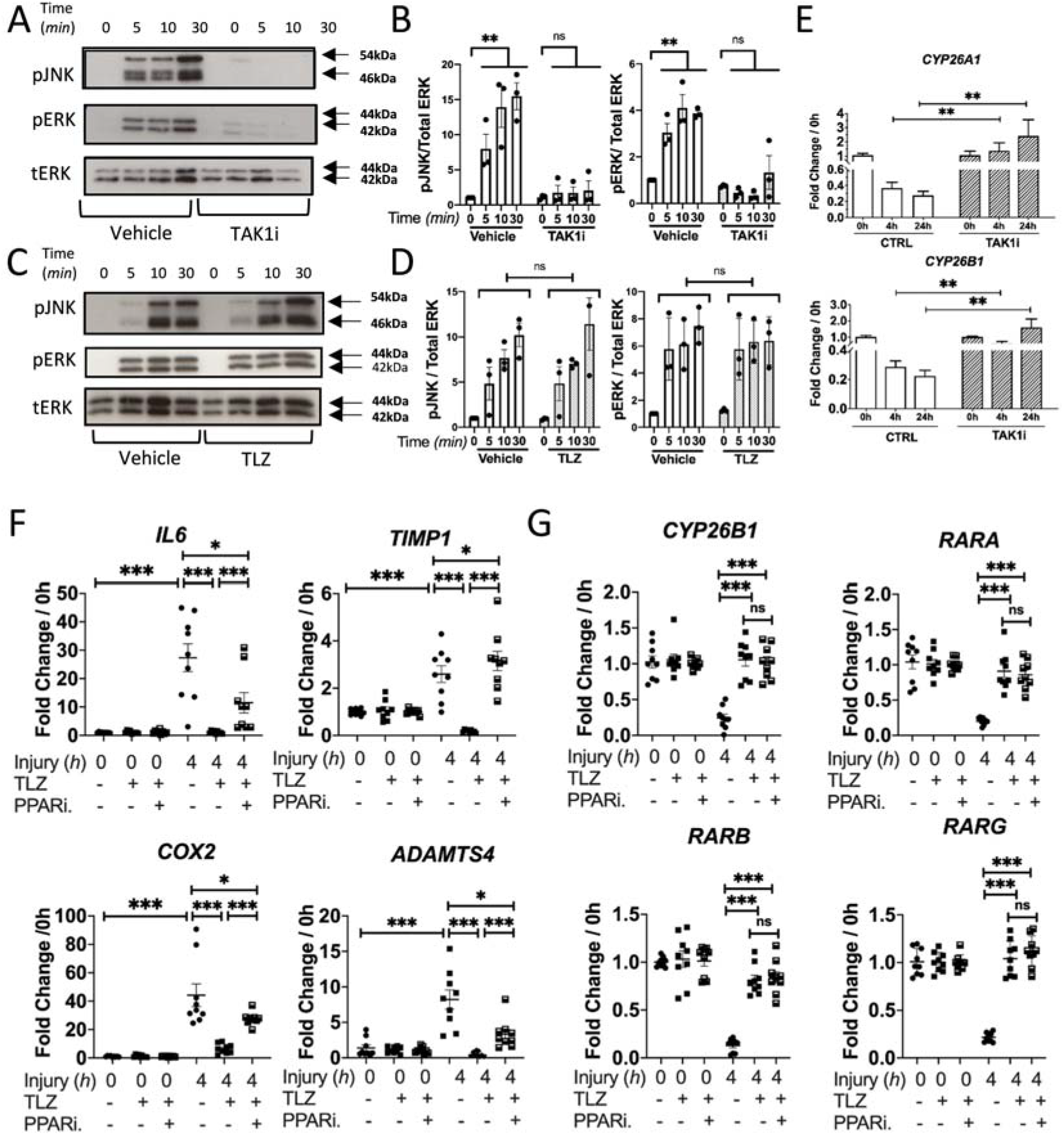
Talarozole does not modulate inflammatory MAPK signalling but suppresses mechanoflammation at the transcriptional level through a PPARγ-dependent manner. Western blot analysis was conducted to investigate the phosphorylation of JNK and ERK in injured porcine cartilage explants in the presence of a TAK-1 inhibitor (5 μM) (TAK1i) (A) or talarozole (5 μM) (TLZ) (B). Porcine MCP joints were pre-injected as described in Fig. 4 but were cultured for 5, 10 or 30 min before lysis. Lysates were immunoblotted for phospho-JNK (p-JNK), phospho-ERK (p-ERK) and total-ERK (t-ERK). Representative immunoblots (A, C), with quantification of three independent experiments (B, D) are shown. (E) Gene regulation of atRA-responsive genes 4h after injury in the presence of TAK1i (mean ± SEM, n=3 by two-way ANOVA). Gene regulation of inflammatory response genes (F), or atRA-responsive genes (G), after pre-injection with a PPARγ inhibitor (10 μM), TLZ (5 μM), vehicle or a combination of inhibitors (mean ± SEM, n=9 individual injected joints). Statistical analysis was performed using Two-way ANOVA with Bonferroni’s post hoc analysis. *p < 0.05, **p < 0.01, and ***p < 0.001.

### Subcutaneous administration of talarozole suppresses mechano-inflammatory genes in articular cartilage early after surgical joint destabilisation

To test the *in vivo* relevance of our findings we next investigated whether systemic delivery of TLZ was able to suppress mechano-inflammatory genes in the articular cartilage *in vivo* after destabilistion of the medial meniscus (DMM). Talarozole was delivered subcutaneously by osmotic minipumps, from 3 days before DMM surgery, and cartilage was harvested after 6h (Fig. 6A). *Cyp26a1* and *Cyp26b1* mRNA increased in TLZ treated mice when normalized to the vehicle contralateral control (Fig. 6B), confirming target engagement of TLZ in cartilage after systemic delivery. Of the inflammatory genes regulated in articular cartilage 6h post joint destabilization, *Il1b, Il6, Adamts4, Mmp3, and Ccl2* were significantly suppressed by TLZ (results normalized to respective 0h control). Two up-regulated genes, *Timp1* and *Cox2,* were unaffected by TLZ (Fig. 6C).

**Fig 6.**
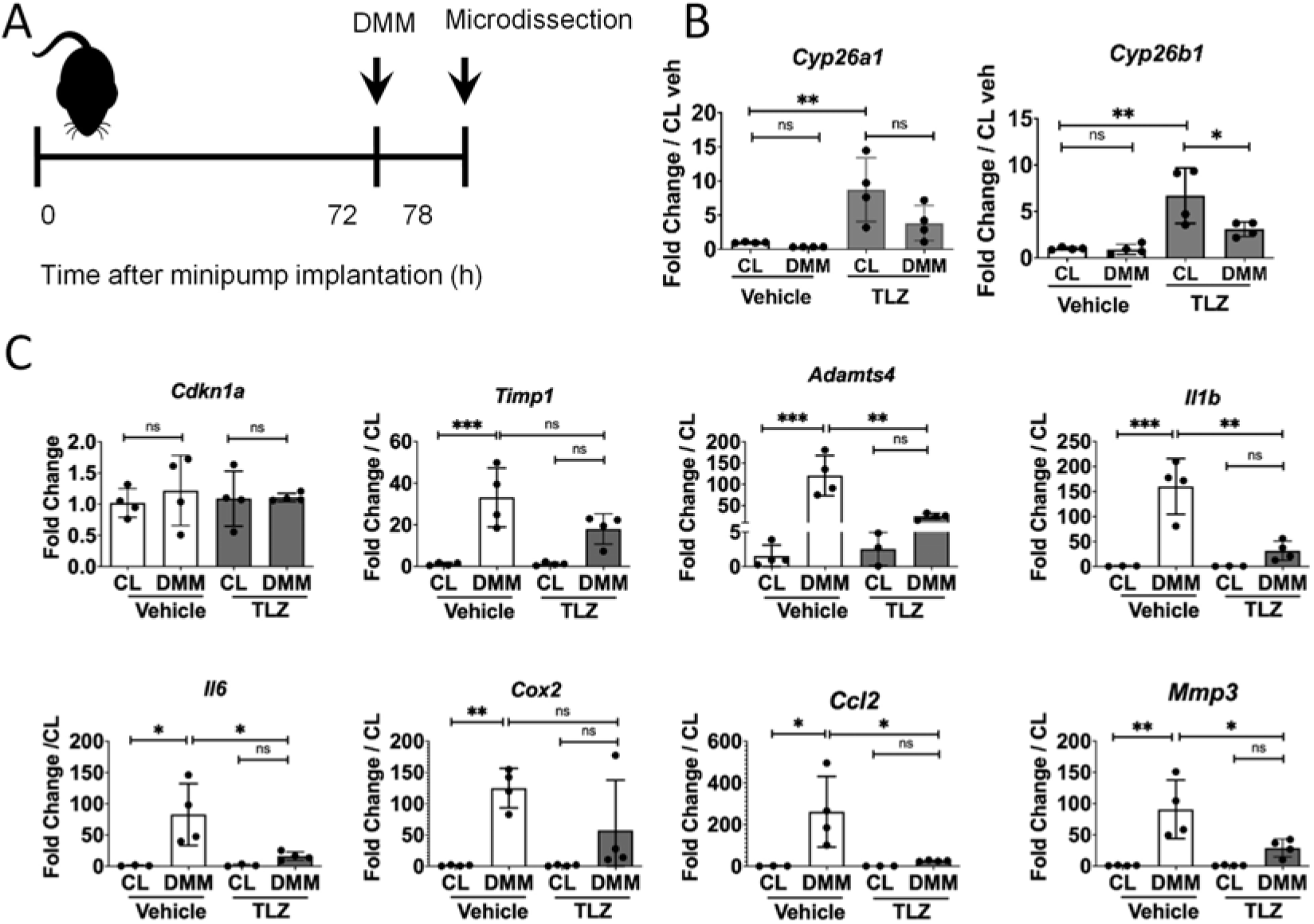
Subcutaneous administration of TLZ by minipump suppressed inflammatory response genes in the articular cartilage 6 hour post joint destabilisation. (A) 10 week old Male C57BL/6 mice were implanted with minipumps containing TLZ (delivering 4 mg/kg) or vehicle. Three days after implantation, destabilisation of the medial meniscus (DMM) was performed on the right knee joint. Six hours post DMM, cartilage was micro-dissected. Cartilage from four mice was pooled for analysis. DMM and contralateral (CL) tissues were snap frozen in RNAlater. (B) Quantitative PCR was performed for atRA-responsive genes and (C) inflammatory response genes (mean ± SEM, n=4). Some genes show fewer than 4 data points due to the undetermined expression level. Results were normalised to *18s* and expressed relative to CL vehicle control (B), or expressed relative to CL for respective treatment group (C). Two-way ANOVA with Bonferroni’s post hoc analysis. *p < 0.05, **p < 0.01, and ***p < 0.001.

### Subcutaneous administration of TLZ attenuates the development of post-traumatic OA

Finally, we investigated whether systemic administration of TLZ could prevent post-traumatic OA in mice. OA was induced by partial meniscectomy (PMX). TLZ was delivered for 23 days (from 2 days prior to surgery to 3 weeks after surgery), and joints harvested at 4 weeks (Fig. 7A). Statistically significant reduction in histological cartilage degradation (OARSI score) (Fig 7B and 7C) and marked reduction in osteophyte size (Fig. 7D) and maturity (Fig. 7E) were observed in the TLZ treated group.

**Fig 7.**
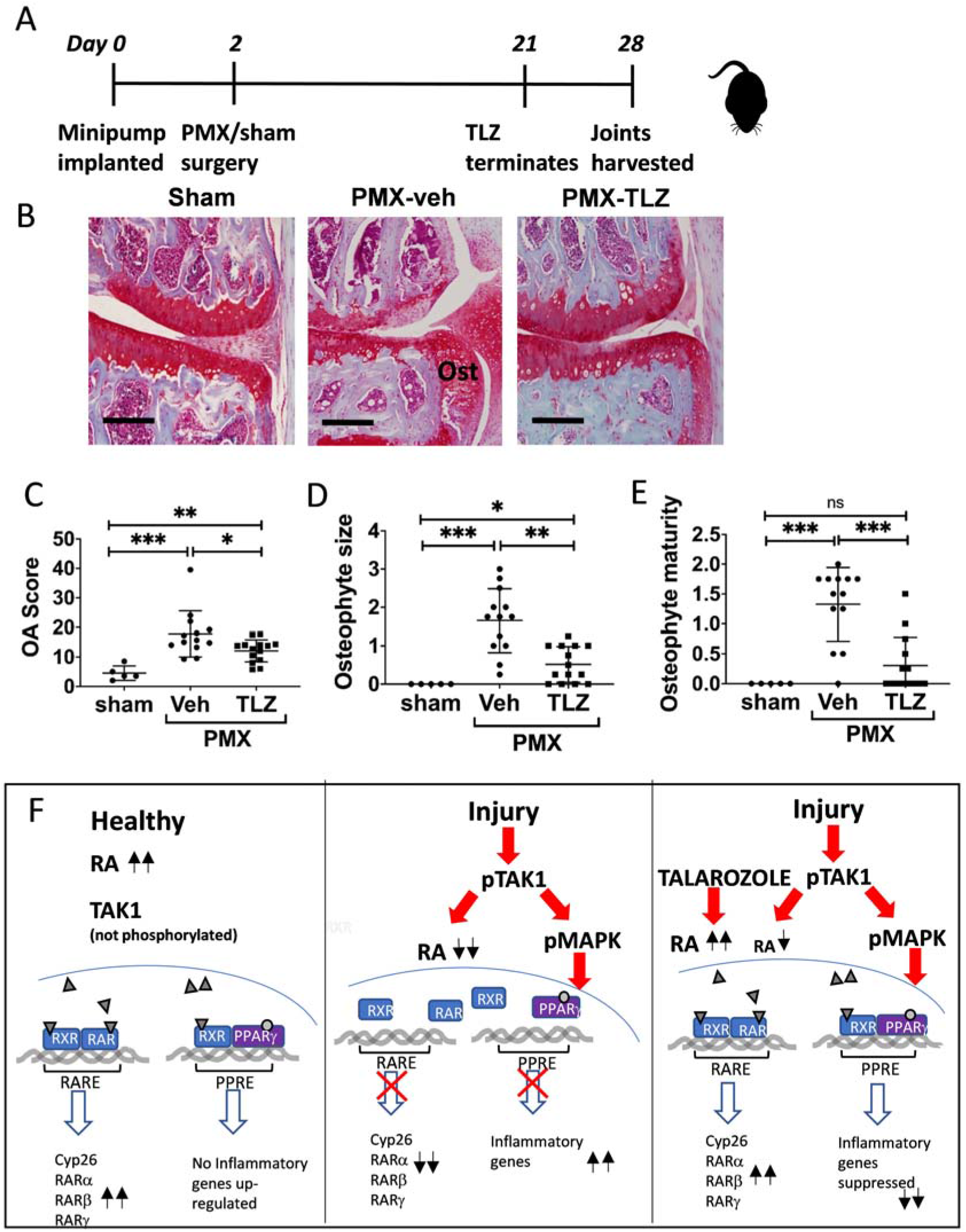
TLZ attenuates the development of post-traumatic OA; suppressing chondropathy and reducing osteophyte formation. (A) Time course for the *in vivo* experiments. 11-week-old male C57BL/6 mice were implanted with minipumps containing TLZ (delivering at 2.5 mg/kg) or vehicle control. Two days after the implantation, PMX or sham surgery was performed on the right knee joint. TLZ was delivered by minipump for three weeks and was left in situ for the duration of the experiment. At four weeks, whole joints were harvested for histological scoring. (B) Representative histological sections at 4 weeks. Histological chondropathy scores (OA score) (C), osteophyte size (D) and osteophyte maturity (E) of sham, vehicle and TLZ treated PMX groups. Scale bar = 200 μm. Sham, n=5; PMX-Ctrl, n=13; PMX-TLZ, n=14. Two mice from (one from each group) were culled early due to a problem with the minipump. Statistical significance was determined by Mann□Whitney U tests. *p < 0.05, **p < 0.01, and ***p < 0.001. Error bars denote mean ± SEM. (F) Schematic of anti-inflammatory role of atRA after cartilage injury. atRA is maintained at high levels in healthy tissue leading to constitutive expression of RARE-dependent genes. After injury, TGFβ-activated kinase 1 (TAK1) is phosphorylated which leads to a drop in atRA and the induction of inflammatory genes. The drop in atRA leads to a reduction in RARE-dependent genes and a loss of PPARγ-mediated suppression of inflammatory genes, which is reversed in the presence of talarozole. Blue triangles – atRA or one of its isomers. Blue circle – PPARγ ligand (multiple endogenous candidates including atRA).

## Discussion

In this study, we confirm that *ALDH1A2* allelic variants, discovered in the Icelandic population, are also associated with hand OA in a large UK cohort. RNA-sequencing of hand OA cartilage, stratified according to *ALDH1A2* genotype and gene dosage, suggested a number of differentially regulated cellular networks and identified atRA as a potential anti-inflammatory molecule in articular cartilage. Maintaining atRA levels at the time of cartilage injury, by blocking its cellular metabolism with talarozole, not only abrogated mechano-inflammatory gene regulation by a mechanism involving PPARγ, but prevented structural joint damage *in vivo* in mice with surgically induced OA.

One striking feature in talarozole treated OA mice was suppression of osteophyte development. These results suggest that atRA is not only suppressing mechanoflammation in articular cartilage but also affecting mechano-sensitive signalling in bone. It may be relevant that PPARγ strongly suppresses osteogenesis (*34*). Florid osteophyte formation and bone expansion in hand OA is often observed in severely affected patients and could explain why *ALDH1A2* variants are associated with symptomatic hand OA rather than large joint disease (*35*). Our results indicate that atRA-dependent genes fall into two principal categories. Those genes that are directly induced by atRA such as *CYP26* (*A1, B1*) and *RAR* (*A, B* and *G*). These genes have classical RAREs in their promoter regions (*17*). The second group are those that are suppressed in an atRA-dependent manner. These appear to be largely dependent on PPARγ suggesting that the RXR-PPAR heterodimer is an important anti-inflammatory regulator in chondrocytes. Anti-inflammatory actions of both atRA and PPARγ have been described previously including by transcriptional activation of IκBα, epigenetic histone modification and direct repression of gene transcription through PPRE binding (*36–39*). Our study highlights that atRA controls the anti-inflammatory effects of PPARγ. This conclusion is consistent with the observation that genetic deletion of PPARγ accelerates surgically induced murine OA (*40*), but PPARγ agonists are not disease modifying (*41*). In essence, we conclude that atRA determines the activity of PPAR-RXR transcriptional regulation and thus controls anti-inflammatory actions of PPARγ (Fig. 7F).

The mechanism by which inflammation is triggered by cartilage mechanical injury remains unclear. In vascularized tissues this involves a breach of the endothelium, exposure of tissue factor and activation of platelets and immune cells (*42*). Cartilage is avascular and so these mechanisms are not relevant. As phosphorylation of MAPKs upon injury is not affected by TLZ, but TAK1 inhibition prevents the drop in atRA-dependent genes, our data suggest that the drop in cellular atRA levels modulates the inflammatory response but is not the primary driver of inflammatory signalling. The cellular mechanisms underlying the activation of MAPKs and the drop in atRA upon injury are currently being pursued in our laboratory.

atRA analogues have been used to increase atRA levels for the treatment of acute promyelocytic leukaemia (*22, 43*), inflammatory disorders such as psoriasis (*44–48*), and inflammatory intestinal diseases (*28, 49*). RAMBAs such as ketoconazole, liarozole, and talarozole offer advantages over atRA analogues and have been used in clinical trials (*21, 22, 50*). Among these, TLZ is the most CYP26-specific inhibitor currently available with IC_50_ of 4-5 nM (*50*). As talarozole has shown therapeutic efficacy in psoriasis and acne vulgaris in humans, with an acceptable safety profile (*21*), our experimental findings make a compelling case for blocking atRA metabolism using a RAMBA to modify disease in OA.

There are a number of recognised limitations in our study. Firstly, we always recognised that we were likely to be underpowered when performing an RNA sequencing exercise on human diseased cartilage samples stratified by polymorphic variants. This was only possible as the polymorphic variants were common in the population (around 50%), in linkage disequilibrium with one another, and the variants conferred a sizeable change in the expression level of *ALDH1A2*. Consequently, we were able to identify a limited number of genes significantly regulated by genotype, but were able to glean more about the biology when we examined *ALDH1A2* gene correlations. This was sufficient to allow us to test some specific hypotheses to verify that atRA was an important anti-inflammatory agent in chondrocytes and amenable to pharmacological interference. Our *in vivo* experiments were substantially restricted by talarozole’s short half-life, requiring it to be delivered continuously through minipump. Some of our mice failed to tolerate the pumps and had to be culled early. This was further complicated by talarozole’s low solubility which limited the total amount of drug that could be carried by the minipump and thus limited the maximum delivery time to three weeks. We were therefore only able to assess the disease modifying ability of talarozole in post-traumatic OA at one early time point. Nonetheless, these results showed significant structure modification with marked effects on osteophyte formation as well as cartilage damage. As the mice were carrying subcutaneous minipumps on their backs, we were unable to assess pain behaviour by incapacitance testing, as per our usual practice. Some of these aspects will be addressed in later studies after incorporation of talarozole into the chow or by manufacturing slow-release preparations.

## Materials and Methods

### Study Design

The aim of this study was to uncover the role of atRA in OA using a combination of human OA tissue taken at the time of hand OA surgery, and normal mouse and porcine articular cartilage subjected to mechanical injury either *in vitro* or *in vivo*. RNAseq of human tissue was always likely to be underpowered. We collected 33 human samples (the largest recorded collection of this sort) of which 19 samples went into the sequencing analysis (12 homozygous for the risk variant and 7 homozygous for the low risk variant). Using these numbers we were able to use PhenomeExpress and Spearman correlation to develop hypotheses that could be tested more robustly in other models. *In vitro* cartilage injury studies are well developed in our laboratory and 3-6 samples is typically what we have used in the past to show robust regulation of genes upon injury. These numbers were empirically increased when using pharmacological inhibitors. Typically, we also increase the concentration of inhibitor (10-fold) when injecting into a joint cavity compared with what we would use *in vitro*. For *in vivo* murine experiments we performed power calculations assuming a 50% reduction in cartilage degradation score using talarozole. The PMX model is more aggressive than DMM (destabilisation of the medial meniscus), that is used by most others in the field, and this allowed us to assess the effect of the inhibitor at an early time point, 4 weeks. All surgery was performed by one operator (JZ). Mice were randomised to vehicle or talarozole pumps, and to sham or PMX surgery. Scoring of histological sections was performed blind by two independent scorers (JZ and LZ). Two mice were excluded from the analysis as they needed to be culled early after surgery due to problems with the minipump. Otherwise, all data were included (outliers were not excluded from any analyses). N numbers used in each experiment are indicated in the respective figures/ legends.

### Reagents and antibodies

Cyp26 inhibitor talarozole (A3853-APE) was obtained from Stratech. All-trans retinoic acid (R2625) was purchased from Sigma-Aldrich. Anti-ALDH1A2 antibody (ab75674) was purchased from Abcam (MA, USA); anti-total ERK (T-ERK) (sc-94) was obtained from Osmotic pumps (2001, 2002) were purchased from Alzet Osmotic pumps (Canada). The kits for RNA extraction were from Qiagen (RNeasy Mini Kit and RNeasy Micro Kit). The reverse transcription kit, PowerUp™ SYBR® green master mix and TaqMan™ universal PCR master mix were from Applied Biosystem (CA, USA). DNAzol™ Reagent (10503027) was from ThermoFisher Scientific. The DNA clean kit (DNA Clean & Concentrator™-5) was from Zymo Research. Anti-phospho-SAPK/JNK antibody (#9251) and anti-phospho-p44/42 MAPK (Erk1/2) antibody (#9101) were obtained from Cell Signaling Technology (MA, USA). The PPARγ inhibitor, GW9662, was from Sigma-Aldrich.

### Human hand OA tissue

UK Biobank has approval from the North West Multi-Centre Research Ethics Committee (11/NW/0382). This study was part of UK Biobank Project ID 22572 (“The Genetics and Epidemiology of Common Hand Conditions”). Trapezia were collected at the time of surgery (from individuals with severe base of thumb OA). Human tissues, generated as part of the procedure, were collected in an approved project within the Oxford Musculoskeletal Biobank. Tissue collection, storage and processing was undertaken following informed written consent and in full compliance with national and institutional ethical requirements, the UK Human Tissue Act, and the Declaration of Helsinki (HTA Licence 12217 and REC reference number 09/H0606/11). The trapezium cartilage was dissected from the bone within 2 hours of surgical removal from the joint and was snap frozen in liquid nitrogen and stored at −80 °C before RNA extraction. In total 33 trapezium samples were collected.

### Human single nucleotide polymorphism genotyping

Human trapezium bone tissue was ground to a powder using Cryo-Cup Grinder (Biospec, Bartlesville), from which genomic DNA (gDNA) was extracted using DNAzol (10503027, ThermoFisher Scientific). Primers were designed based on the 1000 nucleotide sequence adjacent to the target SNP allele, and amplified PCR fragments of the expected sizes were confirmed by Agarose gel electrophoresis. Primers for rs3204689: CATAGGTTTGATTTGTTCCTTCTC (forward), GGTTCGTTCTTATTCTGCCAC (reverse), and for rs4238326: CACACACACCCCAAAACTG (forward) and GGGATAAAGGGAGGAGGAGG (reverse). The purified PCR products were Sanger sequenced by Eurofins Genomic.

### RNA Sequencing

After genotyping, only samples that were homozygous at both rs3204689 and rs4238326 for the wild type (WT) or alternate allele (HO) were processed for RNA-sequencing. (WT n = 8, HO n = 12). Total RNA was extracted using the Qiagen RNeasy Micro Kit according to the manufacturer’s instructions, and its concentration was determined with the NanopDrop 1000 spectrophotometer and RNA quality assessed with the Agilent Bioanalyzer. PolyA-selected sequencing libraries were prepared using the TruSeq protocol (Illumina). Libraries were subject to 75 bp paired-end sequencing (Illumina HiSeq 4000). One sample (WT; OMB0901) was excluded from the analysis due to the low RIN number (5.5). Sequence reads were aligned to the human genome with Hisat2 (version 2.1.0) (*51*) using a “genome_trans” index built from the GRCh38 release of the human genome and Ensembl version 91 annotations (two-pass strategy to discover novel splice sites; with parameters: --dta and --score-min L,0.0,-0.2). Data quality was assessed using pipeline_readqc.py (https://github.com/cgat-developers/cgat-flow/). The average alignment rate was 97.15% (as assessed with picard tools v2.10.9, https://github.com/broadinstitute/picard). Mapped reads were counted using featureCounts (Subread version 1.6.3; Ensembl version 91 annotations; with default parameters) (*52*). Salmon v0.9.1 was used to calculate TPM values (*53*) using a quasi-index (built with Ensembl version 91 annotations and k=31) and gc bias correction (parameter “-- gcBias”).

Differential expression (DE) analysis of WT vs HO samples was performed using Bioconductor package DESeq2 (v1.22.0) (*54*). Prior to analysis, genes were filtered so as to retain n=14,195 protein coding genes expressed at a TPM > 1 in at least three samples of one of the two conditions. Based on exploratory analysis and expert biological knowledge the variables “Tissue from hand” (i.e. whether the tissue was taken from the dominant hand) and “gender” were included as co-variates in the linear model. Outliers counts (n=89 genes) were excluded (Cook’s distance cutoff = 5). Gene set enrichment analysis with FGSEA was performed using the n=14,106 genes tested for differential expression by DESeq2 (i.e. after independent filtering) using the multilevel procedure with GO Biological Process (BP), Cellular Component (CC) and Molecular Function (MF) and KEGG pathway annotations (*55*). To help remove redundant pathways we applied the FGSEA ‘collapsePathways’ function (results provided in Supplemental Data File S2). The metadata used for the differential expression analysis are provided in Supplementary Data File S5.

The PhenomeExpress analysis was performed using the Cytoscape PhenomeScape app (*29*). The Human network was loaded, and the set of genes tested for differential expression with DESeq2 analysis (n=14,106 genes, supplementary Data files S4) was used as the input for the targeted Osteoarthritis (HP:0002758) phenotype analysis.

For the *ALDH1A2* gene expression level correlation analysis, genes were not filtered by biotype. Lowly expressed genes were removed (average log CPM values < −1; as computed using the ‘aveLogCPM’ edgeR function (*56*)), and counts for the remaining n=17,348 genes normalized with DESeq2 and subjected to variance stabilizing transformation prior to computation of the Spearman correlations. FGSEA analysis was performed as before with the n=17,348 genes that passed the initial expression filter using Spearman’s rho as the ranking metric (full results provided in Supplemental data file S4) (*55*).

### Cartilage injury and culture

Porcine cartilage: articular cartilage was dissected from metacarpophalangeal (MCP) joints from trotters (forelimb) of freshly slaughtered 3-6 months old male pigs from a local supplier. Trotters were first immersed in 2% Virkon for decontamination, and then equilibrated at 37 0C for one hour. 1 ml of serum free Dulbecco’s modified Eagle’s medium (DMEM) with or without inhibitor (Cyp26 inhibitor (talarozole) (5 μM), GW9662 (PPARγ inhibitor) (10 μM), or TAK1 inhibitor (5 μM)), or vehicle was injected into the MCP joint. Trotters were incubated for another hour at 37 °C. The joint was then opened and cartilage was rapidly dissected from the articular surface either directly onto dry ice (0h) or was cut into small pieces as described previously (*57*) and cultured for the indicated times in medium with or without inhibitor. Tissue extracts were generated using radioimmunoprecipitation assay (RIPA) buffer containing phosphatase inhibitors (45min on ice) for western blotting or Trizol for qPCR.

### RNA extraction and real-time quantitative reverse transcription–polymerase chain reaction (qPCR)

Total RNA extraction from porcine cells, cartilage explants and murine tissue was performed using a Trizol and Qiagen RNeasy Mini Kit according to the manufacturer’s instructions. Complementary DNA (cDNA) was synthesized from RNA using a high capacity cDNA Reverse Transcription Kit (Applied Biosystems). Primers for porcine genes were designed and listed in Table S3, and real-time PCR was performed with the ViiA™ 7 system (Applied Biosystems), using SYBR green Master-Mix (Applied Biosystems) according to the manufacturer’s instructions. Expression of the respective genes was normalized to that of *18s* as an internal control, using the ΔΔCt method. Human trapezial cartilage was ground to a powder using Cryo-Cup Grinder (Biospec, Bartlesville) and then RNA was extracted using Qiagen RNeasy Micro Kit.

### TaqMan Low-Density Array (TLDA) microfluidic cards

Complementary DNA (cDNA) was generated from RNA using a High Capacity cDNA kit (Applied Biosystems) according to the manufacturer’s instructions. TLDA microfluidic cards were custom designed to order from Applied Biosystems. All thermocycling was carried out on the ViiA™ 7 system (Applied Biosystems). 50 μl cDNA template in nuclease-free water and 50 μl of TaqMan Universal PCR Master Mix (Applied Biosystems) were mixed and added to each TLDA loading port. The TLDA card was then centrifuged and sealed. Ct values were obtained after manually choosing the Ct threshold. Expression of the respective genes was normalized to that of *18s* as an internal control, using the 2^−ΔΔCt^ method. The Taqman probes used for human and mouse genes were purchased from Applied Biosystems and listed in Table S5.

### Western blotting

Porcine cartilage explant lysates were resolved by 10% sodium dodecyl sulfate– polyacrylamide gel electrophoresis, and immunoblotted for ALDH1A2 or for phosphorylated proteins p-JNK, p-ERK and total ERK. Quantification of three separate experiments was performed using FIJI-App, ImageJ-win64 Gel Analysis program, normalised to total ERK1/2.

### Animals

Animal experiments were carried out under ethical approval in agreement with local policy. 4-5 mice per cage were housed in standard individually ventilated cages under a 12 hour light/dark cycle. 4 or 10-week-old C57BL/6 male mice were obtained from Charles Rivers, UK laboratory.

Talarozole *in vivo* experiment: 11-week-old mice received talarozole via pump (mini-osmotic pump, Alzet, model 2001 and 2002) for a period of 3 days or 4 weeks. Talarozole was dissolved in PEG 300 and DMSO (50:50) to a daily dosage of 4 mg/kg/day (2001, 3 days) or 2.5 mg/kg/day (2002, 4 weeks). Mice undergoing surgery were anesthetized by inhalation of isoflurane (3% induction and 1.5–2% maintenance) in 1.5–2 liters/minute oxygen. All animals received a subcutaneous injection of buprenorphine (Vetergesic; Alstoe Animal Health) immediately before surgery. Mice were fully mobile within 3–5 minutes after withdrawal of isoflurane. A small incision was made on the right flank and a mini pump was inserted subcutaneously and the wound was closed by suture. For the short time experiment, destabilization of the medial meniscus (DMM) surgery was conducted on the right leg of mice as previously described (16). 6 hours post DMM, mice were culled by CO_2_, and cartilage from DMM joint and contralateral joint was collected separately by microdissection as described previously. For the longer time experiment, partial meniscoectomy was carried out two days after the minipump implantation. These mice were culled four weeks post the implantation of the minipump and right joints were harvested.

### Immunohistochemistry

Paraformaldehyde-fixed joints from mice 4 weeks postsurgery (PMX or sham) were decalcified in EDTA for 2 weeks and then embedded in paraffin. Joints were sectioned coronally and stained with Safranin O, FastGreen and hematoxylin for microscopic inspection and histological scoring. Histological analyses of the knee joints was done, according to a standard system, by 2 observers under blinded conditions, and a summed score (sum of the 3 highest total section scores for any given joint, minimum of 10 sections per joint, 80 μm apart) was obtained. Osteophytes were scored by size (0-3) and maturity (0-3) as previously described. ^(*58*)^

### Statistical analysis

Data for replicate experiments are expressed as the mean ± SEM. Analysis was performed using Prism 7 software (GraphPad). Student’s t-tests were used to establish statistical significance between two groups. One-way analysis of variance (ANOVA) was used to compare multiple groups. Two way ANOVA with Bonferroni’s post test was performed for multiple comparisons. Mann-Whitney U test was used to analyse PMX-Ctrl and PMX-TLZ histological pathology scores. p values less than 0.05 were considered statistically significant.

## Supplementary Materials

### Materials and methods

Table S1. Characteristics of individuals for each hand OA sample included in the study. Table S2. Summary of PhenomeExpress Networks in Fig. 1.

Table S3. Primers and Taqman probes used for Real-Time PCR.

Fig. S1. Differentially expressed genes and enriched biological pathways in WT vs HO samples.

Fig. S2. Pathways enriched in genes showing correlation in expression with *ALDH1A2*. Fig. S3. atRA-responsive genes in porcine chondrocytes.

Fig. S4. Reciprocal relationship between atRA-responsive and inflammatory genes following murine cartilage injury.

Fig. S5. Exogenous atRA or talarozole supresses IL-1 induced inflammatory response genes in isolated porcine chondrocytes.

Data file S1. Differential expression analysis results, WT vs HO.

Data file S2. Gene set enrichment analysis (FGSEA) against Ontology and KEGG pathways using differential expression analysis results, WT vs HO.

Data file S3. Significant Spearman Correlation Coefficients of expression of ALDH1A2 in hand OA cartilage, relating to Fig. 2.

Data file S4. Gene set enrichment analysis (FGSEA) against Ontology and KEGG pathways using gene list of Spearman Correlation analysis, WT vs HO.

Data file S5. Metadata for Human RNA seq.

## Acknowledgements

Tissue samples and/or data were obtained from the Oxford Musculoskeletal Biobank and were collected with informed donor consent in full compliance with national and institutional ethical requirements, the UK Human Tissue Act, and the Declaration of Helsinki (HTA Licence 12217 and Oxford REC C 09/H0606/11). We thank the Oxford Hand Surgery Team who include: Nick Riley, Michelle Spiteri, Ian McNab, Christopher Little, Lucy Cogswell, Paul Critchley, Henk Giele, Rebecca Shirley and our centre tissue coordinators Louise Hill and Katherine Groves for facilitating collection of tissues. We thank Dr. Mino Medghalchi and Mr. Albertino Bonifacio for their technical support with the minipump implantation. We thank colleagues from the Kennedy histology lab Ida Parisi and Grace Wilson for their kind help with the immunohistochemistry staining. We thank Moustafa Attar for his help with the RNA-seq sample processing.

## Funding

Centre for OA Pathogenesis Versus Arthritis (grant numbers 20205 and 21621). National Institute for Health Research (NIHR) Oxford Biomedical Research Centre (BRC) Wellcome Trust Intermediate Fellowship to DF (WT097152). UKRI Future Leaders Fellowship to FEW (MR/S016538/1).

## Author contributions

Study design: LZ, SS and TLV

Method development, data acquisition and analysis: LZ, PK, LK, JMZ, AC, HI, HM.

Recruitment of patients, tissue acquisition and clinical governance: DF, FEW, MDG, the Oxford Hand Surgeons.

Replication of SNP analysis in UK Biobank: DF, AW, MN Writing of manuscript: LZ, TLV.

All authors critically reviewed the final draft.

Disclaimer: The views expressed are those of the author(s) and not necessarily those of the NHS, the NIHR or the Department of Health.

## Notes

### Competing Interest Statement

The authors have declared no competing interest.

### Summary of Updates

Figure 1 and 2 revised. RNAseq analysis revised. Supplemental files updated.

