## Supplementary material for "Variants in *ALDH1A2* reveal an anti-inflammatory role for retinoic acid and a new class of disease-modifying drugs in osteoarthritis": supplemetary

**Supplementary Methods:**

**Materials and Methods**

**UK Biobank replication study**

The analysis was performed after approval from UK Biobank (application number 22572). Hand OA cases were selected from UK Biobank participants on the basis of the following diagnostic codes under either Primary or Secondary ICD-10 diagnoses: M18 (Arthrosis of first carpometacarpal joint + all sub-codes), M15.1 (Heberden’s nodes with arthropathy), and M15.2 (Bouchard’s nodes with arthropathy). The genotyping, QC and imputation methodology employed by UK Biobank are described in detail elsewhere (*59*), and full details of our SNP- and sample-level quality control (QC) pipeline have been previously described (*60*). Briefly, SNP-level QC reduced the number of SNPs from 784,256 to 547,011, and sample-level QC reduced the cohort from 488,377 to 401,656 individuals of white British ancestry confirmed on principal component analysis. There were 2,140 hand OA cases post-QC.

Matching was performed using R V8.3.1, and we used the R package MatchIt (*61*) to match hand OA cases to controls. Controls were selected from the non-hand OA UK Biobank population, and were matched to cases 1:5 based on the following parameters, using the “*nearest*” matching method: sex, year of birth, genotyping platform (UK BiLEVE Axiom Array or UK Biobank Axiom Array), and UK Biobank recruitment centre. 2,140 hand OA cases were matched to 10,700 controls. Genome-wide association testing was undertaken across 547,011 common-frequency genotyped SNPs (minor allele frequency (MAF) ≥ 0.01) and ~8.4 million imputed SNPs (MAF ≥ 0.01, Info Score ≥ 0.9) using a linear mixed non-infinitesimal model implemented in BOLT-LMM v2.3 (*62*) to account for population structure and relatedness. We extracted association summary statistics (including odds ratios and p-values) for the two SNPs of interest (rs3204689 and rs4238326), and considered a statistically significant association to be less than the Bonferroni corrected p-value for two tests (0.05/2 = 0.025).

**Porcine chondrocyte isolation and culture**

Freshly dissected porcine cartilage was chopped finely and placed in DMEM with 10% fetal calf serum (Invitrogen) and 1.0 mg/ml type II collagenase (Roche). Tissue was incubated at 37^0^C overnight (18-24h). Cells were plated at a density of 2 million cells in each well of 6-well tissue culture plates in complete DMEM containing 10% fetal bovine serum, 1% penicillin/streptomycin, and amphotericin. Prior to stimulation with atRA, IL-1, talarozole, or vehicle, cells were serum starved for 24h.

**Cartilage injury and culture**

Mouse cartilage: 5-6 week-old mice (C57BL/6) were culled by CO_2_. The hip joint was exposed by blunt dissection and the femoral cap was avulsed using forceps, as described previously (7). Hip cartilage was snap frozen (0h) or cultured for 4h post avulsion in serum free medium. Four hips were pooled for one data point. 16 mice were used in total. All dissected cartilage was stored at -80 ^0^C until extraction of total RNA.

**Table S1. Characteristics of individuals for each hand OA sample included in the study.**

| **Sample ID** | **Age** | **Gender** | **Genotype** | |
| --- | --- | --- | --- | --- |
|  |  |  | **SNP rs4238326*** | **SNP rs3204689^#^** |
|  |  |  | **T/C** | **G/C** |
| **OMB15/0198** | 60 | M | WT | WT |
| **OMB15/0230** | 54 | M | WT | WT |
| **OMB15/0383** | 58 | M | WT | WT |
| **OMB15/0252** | 69 | F | WT | WT |
| **OMB0022** | 72 | F | WT | WT |
| **OMB0862** | 54 | F | WT | WT |
| **OMB0901** | 75 | F | WT | WT |
| **OMB0894** | 54 | F | WT | WT |
| **OMB15/0173** | 58 | F | HO | HO |
| **OMB15/0206** | 60 | F | HO | HO |
| **OMB15/0319** | 69 | F | HO | HO |
| **OMB15/0373** | 52 | F | HO | HO |
| **OMB15/0384** | 69 | F | HO | HO |
| **OMB15/0393** | 54 | F | HO | HO |
| **OMB0052** | 63 | F | HO | HO |
| **OMB0481** | 71 | F | HO | HO |
| **OMB0861** | 66 | M | HO | HO |
| **OMB0866** | 83 | M | HO | HO |
| **OMB0877** | 75 | F | HO | HO |
| **OMB0885** | 64 | F | HO | HO |
| **OMB15/0185** | 56 | F | WT | HO |
| **OMB15/0315** | 50 | F | WT | HET |
| **OMB0023** | 57 | M | WT | HET |
| **OMB15/0244** | 50 | M | HO | HET |
| **OMB0027** | 58 | M | HET | HET |
| **OMB0061** | 75 | F | HET | HET |
| **OMB0087** | 65 | M | HET | HET |
| **OMB0140** | 68 | F | HET | HET |
| **OMB0316** | 62 | F | HET | HET |
| **OMB15/0275** | 67 | F | HET | HET |
| **OMB15/0380** | 50 | F | HET | HET |
| **OMB0529** | 54 | F | HET | HET |
| **OMB0876** | 65 | F | HET | HET |

^*^For SNP rs4238326, WT represents homozygocity for the ancestral T allele, HO represents homozygocity for the alternate C allele, and HET represents heterozygocity.

^#^For SNP rs3204689, WT represents homozygocity for the ancestral G allele, HO represents homozygocity for the alternate C allele, and HET represents heterozygocity.

**Table S2. Summary of PhenomeExpress Networks in Figure 1.** Differentially regulated sub-networks related to OA phenotypes. The size, empirical p-value and function is indicated for each network.

| Network number | Top GO biological process | Size | Empirical P-value |
| --- | --- | --- | --- |
| 1 | innate immune response | 8 | 0.03 |
| 2 | cellular response to drug | 32 | 0.015 |
| 3 | positive regulation of osteoblast differentiation | 46 | 0.001 |
| 4 | innate immune response_1 | 14 | 0.014 |
| 5 | cyclooxygenase pathway | 11 | 0.009 |
| 6 | positive regulation of canonical Wnt signaling pathway | 4 | 0.012 |
| 7 | positive regulation of tyrosine phosphorylation of Stat3 protein | 6 | 0.027 |
| 8 | nucleotide-binding domain, leucine rich repeat containing receptor signaling pathway | 8 | 0.015 |
| 9 | extracellular matrix organization | 8 | 0.017 |
| 10 | anterior/posterior pattern specification | 11 | 0.008 |
| 11 | extracellular matrix organization_1 | 10 | 0.022 |
| 12 | bone mineralization | 9 | 0.048 |
| 13 | phosphatidylinositol-mediated signaling | 10 | 0.014 |
| 14 | cholesterol biosynthetic process | 24 | 0.001 |
| 15 | NADP metabolic process | 4 | 0.005 |
| 16 | anterior/posterior axon guidance | 5 | 0.019 |
| 17 | chemokine-mediated signaling pathway | 11 | 0.002 |
| 18 | endocardial cushion to mesenchymal transition involved in heart valve formation | 5 | 0.014 |
| 19 | cell division | 7 | 0.011 |
| 20 | chemokine-mediated signaling pathway_1 | 22 | 0.014 |
| 21 | complement activation | 16 | 0.006 |
| 22 | extracellular matrix organization_2 | 15 | 0.008 |
| 23 | osteoblast development | 6 | 0.015 |
| 24 | positive regulation of cAMP biosynthetic process | 7 | 0.031 |
| 25 | hemoglobin biosynthetic process | 4 | 0.001 |
| 26 | cellular nitrogen compound metabolic process | 4 | 0.016 |
| 27 | regulation of mitotic nuclear division | 5 | 0.037 |

**Table S3. Primers and Taqman proves used for Real-Time PCR.**

|  | Gene ID | Description | Pig | | Mouse Gene Probe ID | Human Gene Probe ID |
| --- | --- | --- | --- | --- | --- | --- |
|  |  |  | Forward Primer | Reverse Primer |  |  |
| **Reference Gene** | *18s* | eukaryotic 18S rRNA | 5'-TGCAGAATCCTCGCCAATACA-3' | 5'-AGTCGCTCCAAGTCTTCACG-3' | Hs99999901s1 | Hs99999901s1 |
|  | *ALDH1A2* | aldehyde dehydrogenase 1 family member A2 | ***** | | Mm00501306m1 | Hs00180254m1 |
| **atRA Responsive Genes** | *CYP26A1* | cytochrome P450 family 26 subfamily A member 1 | 5'-GCAGCCACATCTCTCATTACTTA-3' | 5'-CTTCAGCTCCTCTCGCACT-3' | Mm00514486m1 | Hs00175627m1 |
|  | *CYP26B1* | cytochrome P450 family 26 subfamily B member 1 | 5'-GCACGGCAGATCCTACAGA-3' | 5'-TGTCCAGAGCATCCGAGTAG-3' | Mm00558507m1 | Hs01011223m1 |
|  | *RARα* | retinoic acid receptor alpha | 5'-CGGAACAAGAAGAAGAAGGAGG-3' | 5'-CCAGAGGTCAATGTCCAGAGA-3' | Mm01296312m1 | Hs00940446m1 |
|  | *RARβ* | retinoic acid receptor beta | 5'-CCGACCTTGTGTTCACCTTC-3' | 5'-CCGTCTCTGTGTCATCCATC-3' | Mm01319677m1 | Hs00977140m1 |
|  | *RARγ* | retinoic acid receptor gamma | 5'-GTCCTCTGGCTACCACTACG-3' | 5'-CACAGCTTCCTTGGACATGC-3' | Mm00441091m1 | Hs01559234m1 |
| **Inflammatory Response Genes** | *ADAMTS4* | ADAM metallopeptidase with thrombospondin type 1 motif 4 | 5'-ACACGCCTCCGATACAGCTT-3' | 5'-GTAGAACGTGGCGTTGAAGGA-3' | Mm00556068m1 | Hs00192708m1 |
|  | *IL6* | interleukin 6 | 5'-ATGCTTCCAATCTGGGTTCAA-3' | 5'-CACAAGACCGGTGGTGATTCT-3' | Mm00446190m1 | Hs00174131m1 |
|  | *COX2* | prostaglandin-endoperoxide synthase 2 | 5'-CCGACAGCCAAAGACACTCA-3' | 5'-CGGAGGTGTTCAGGAGTGTGA-3' | Mm00478374m1 | Hs00153133m1 |
|  | *CCL2* | C-C motif chemokine ligand 2 | 5'-TGTGCCTGCTGCTCACTG-3' | 5'-GCAGCAGGTGACTGGAGAAT-3' | Mm00441242m1 | Hs00234140m1 |
|  | *TIMP1* | TIMP metallopeptidase inhibitor 1 | 5'-GAGCCCCAGAGTTCAACCAGAC-3' | 5'-GGCGGGGGCGTAGATGA-3' | Mm00441818m1 | Hs00171558m1 |
|  | *MMP3* | matrix metallopeptidase 3 | 5'-GGAGTTCCTGATGTTGGTTACTTC-3' | 5'-CAAAACTTTTCCAGGTCCGTCAAA-3' | Mm00440295m1 | Hs00968305m1 |
|  | *CDKN1A* | cyclin Dependent Kinase Inhibitor 1A | 5'-CTCCCAGGGCAGGAAACG-3' | 5'-TTGTTTCCAGCAGGACAAGG-3' | Mm00432448m1 | Hs00355782m1 |
|  | *IL1b* | interleukin 1 beta | 5'-GAGGCAGTGAAATTTGACATGG-3' | 5'-GGCAATGAACAACTTTGGATGGG-3' | Mm00434228m1 | Hs01555410m1 |

* unable to design functioning primers

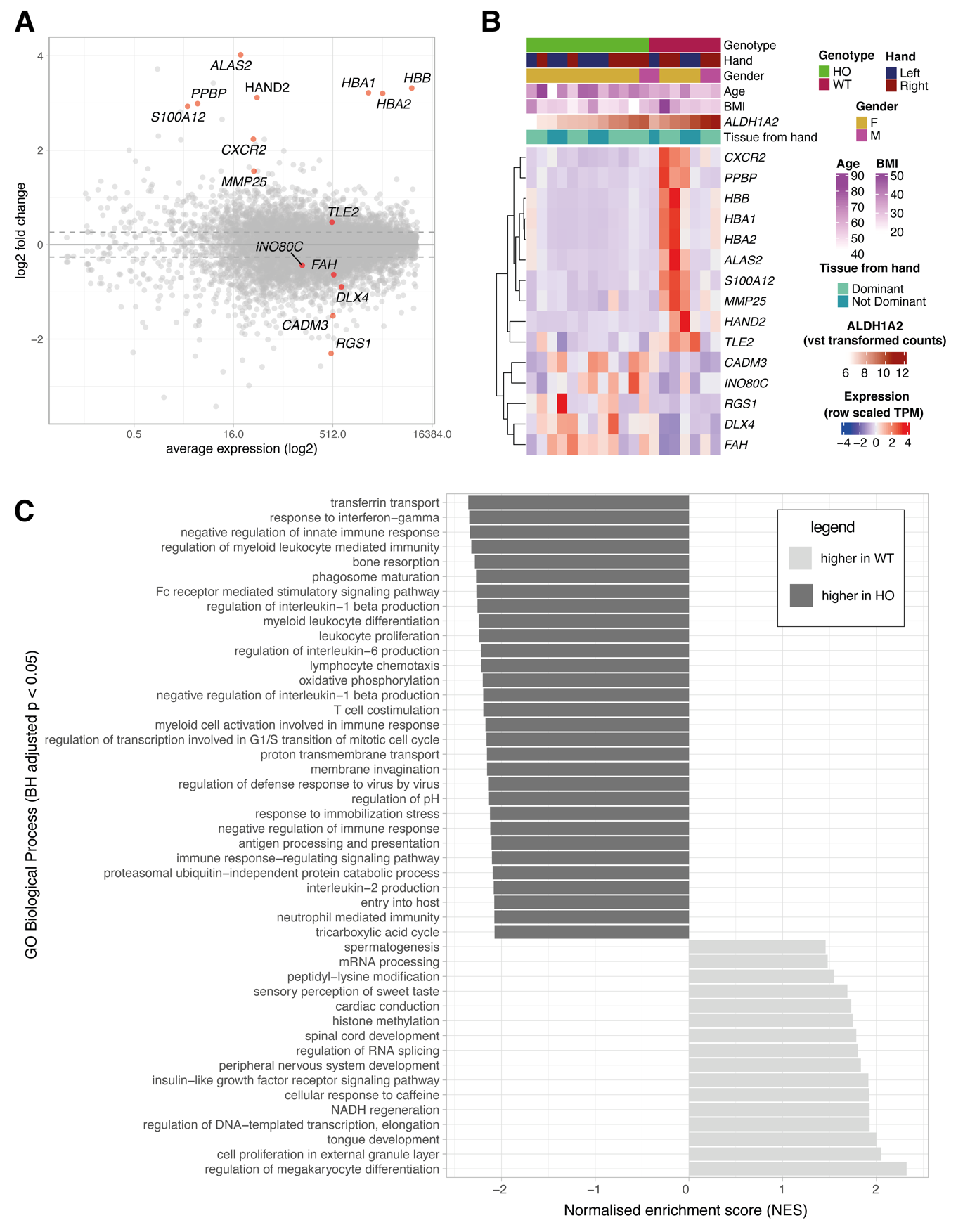

**Fig. S1. Differentially expressed genes and enriched biological pathways in WT vs HO samples.** (A) MA and volcano plot of the n=15 significant differentially expressed genes (DESeq2 analysis, BH adjusted p values < 0.1). Positive fold change values show higher expression in the WT samples. (B) Heatmap of the expression of the n=15 significant differentially expressed genes. (C) Top GO Biological Processes found to be differentially regulated between the WT and HO samples (FGSEA analysis).

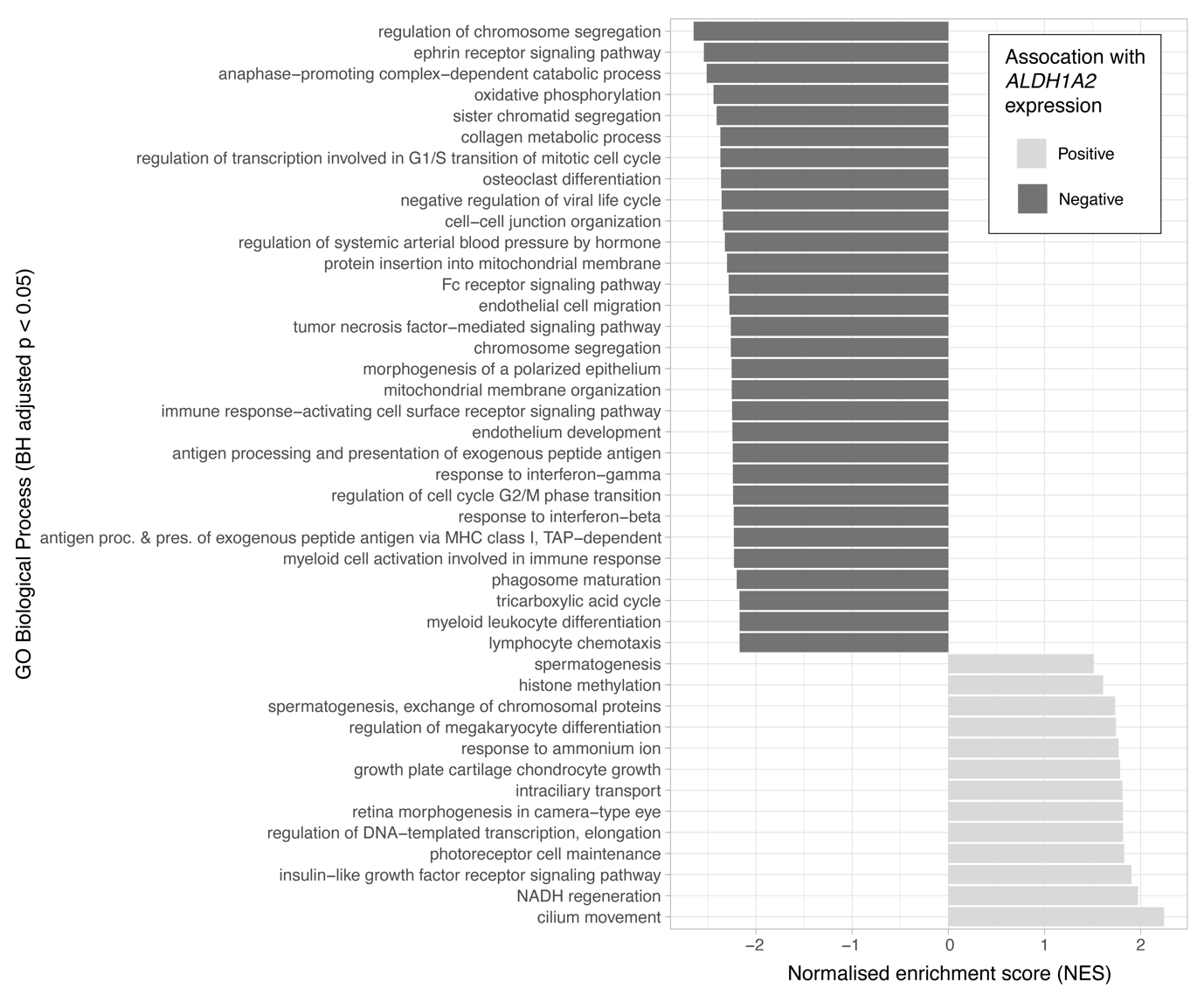

**Fig S2. Pathways enriched in genes that show correlation in expression with *ALDH1A2***. Top GO Biological Processes found to be enriched amongst genes correlated in expression with *ALDH1A2* (FGSEA analysis).

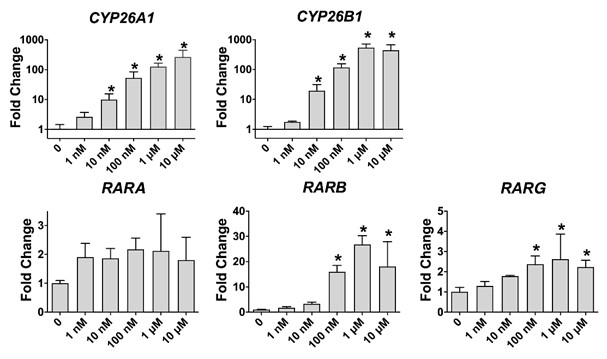

**Fig. S3. atRA-responsive genes in porcine chondrocytes**. Primary porcine chondrocytes were stimulated with atRA at the indicated concentrations or vehicle (0) for 4h. Total RNA was extracted and reverse-transcribed, and the expression levels of atRA-responsive genes were analyzed by qPCR. Fold changes were normalised to *18s* and expressed relative to vehicle (0) (mean ± SEM, n = 3). *p<0.05, *p<0.01, and ***p< 0.001 compared with vehicle (0) by one-way ANOVA.

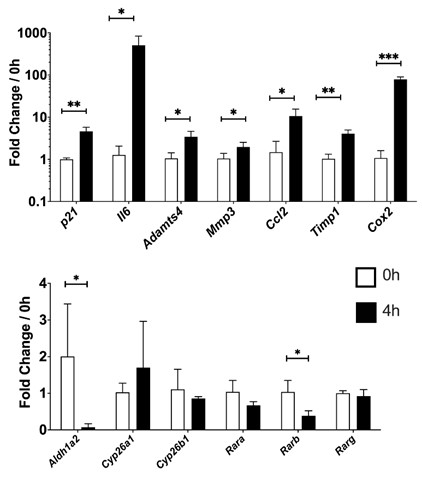

**Fig. S4. Reciprocal relationship between atRA-responsive and inflammatory response genes following murine cartilage injury.** The cartilaginous femoral cap was avulsed using forceps from 5-6 week-old mice (C57BL/6). Hip cartilage was snap frozen (0h) or cultured for 4h post avulsion in serum free medium. Four hips were pooled for one data point. 16 mice were used in total. Selective atRA-responsive genes (A) and inflammatory genes (B) were validated by qPCR. Results were normalised to *18s* and expressed relative to 0h (mean ± SEM, n = 4 experimental replicates for each time point), *p<0.05, **p<0.01 and ***p< 0.001 compared with 0h *by multiple t-test.*

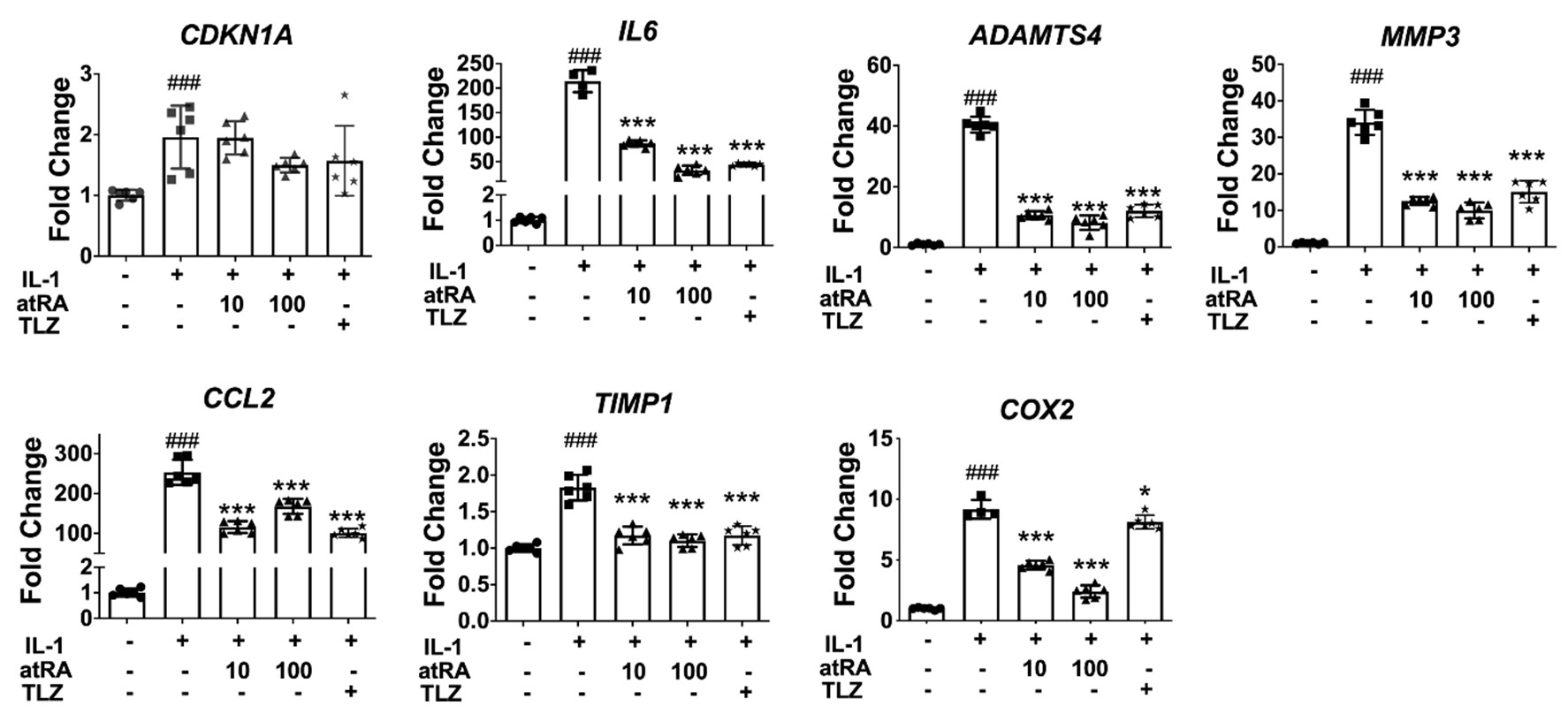

**Fig. S5. Exogenous atRA or talarozole supress IL-1 induced inflammatory response genes in isolated porcine chondrocytes.** Primary porcine chondrocytes were treated with 20 ng/ml IL-1 in combination with different concentrations of atRA or talarozole (500 nM). qPCR was performed for select inflammatory response genes, normalised to *18s* and expressed relative to the untreated control (mean ± SEM, n = 6). ^###^p < 0.001, IL-1 group compared with vehicle control. Statistical analysis was performed using one-way ANOVA with Tukey’s post hoc analysis. *p < 0.05, **p < 0.01, and ***p < 0.001 all other groups compared with IL1.
